# Grass Expression Atlas: an RNA-seq-based expression resource for grass species

**DOI:** 10.64898/2026.03.13.711518

**Authors:** Kota Kambara, Qi Chen, Daisuke Tsugama

## Abstract

Grass Expression Atlas (GExA) is an interactive web-based resource for rapid exploration of gene expression across diverse tissues, developmental stages, and conditions in grass species. GExA integrates publicly available RNA sequencing (RNA-seq) datasets for four millets: pearl millet (*Cenchrus americanus*), foxtail millet (*Setaria italica*), proso millet (*Panicum miliaceum*), and finger millet (*Eleusine coracana*), and includes barley (*Hordeum vulgare*) and sorghum (*Sorghum bicolor*) as reference species. Datasets were processed using a unified processing workflow to generate expression values in transcripts per million (TPM). The current release comprises 4,673 samples from 442 BioProjects, including 987 pearl millet samples and 2,216 foxtail millet samples, and is provided through a user-friendly web interface. GExA is designed for scalable expansion to additional species via the pipeline used in this study. GExA is freely available at https://webpark2116.sakura.ne.jp/RNADB.

## Introduction

Millets are a diverse group of small-seeded grasses cultivated worldwide as important cereal and forage crops. They are gaining increasing attention because several species, including pearl millet (*Cenchrus americanus*), are valued for their nutritional quality and tolerance to abiotic stresses. However, despite their biological and agricultural importance, public bioinformatics infrastructure and functional genomic resources for millets remain limited (Ghatak et al. 2025). Therefore, establishing and strengthening public bioinformatics infrastructure for millets remains an important priority.

High-throughput RNA sequencing (RNA-seq) is widely used to comprehensively and quantitatively profile transcriptomes across tissues, developmental stages, and environmental conditions. For model plants and other crops, web-based gene expression databases have been established by collecting, reanalyzing, and integrating transcriptome datasets generated by multiple independent projects. For example, RiceXPro provides comprehensive expression profiles across organs/tissues and developmental stages in rice (Sato et al. 2011). In barley and sorghum, PlantNexus integrates RNA-seq datasets from multiple projects and offers a global gene coexpression network (GCN) resource (Zhou et al. 2022). In addition, the Plant Public RNA-seq Database compiles and integrates RNA-seq data from more than 45,000 samples across six plant species including rice (*Oryza sativa*) and *Arabidopsis thaliana*, enabling cross-condition and cross-tissue comparisons of gene expression (Yu et al. 2022).

In millets, although publicly available transcriptome datasets have increased (e.g. Lou et al. 2024 and Qazi et al. 2025), user-friendly resources that systematically integrate and reanalyze data across multiple projects remain limited. For pearl millet in particular, no publicly accessible RNA-seq database has been available. Although a previous study reported the collection and processing of pearl millet RNA-seq datasets (Kambara et al. 2024), that resource did not provide a web interface for flexible exploratory analysis. As a result, the practical reuse, comparative exploration, and long-term updatability of public pearl millet transcriptome data remained limited. Similar needs exist for other millets. Although plant RNA-seq resources such as PlantExp (Liu et al., 2023) provide access to public transcriptome data across multiple millet species including foxtail millet (*Setaria italica*), readily usable resources that support integrated exploration of millet gene expression across tissues, developmental stages, cultivars, and treatments remain limited. In particular, tools that enable flexible querying, visualization across projects and biological contexts, and annotation-based exploratory searches would further enhance the usability of public millet transcriptome datasets. These needs motivated the development of an expression resource designed to support efficient exploration and reuse of RNA-seq data across millet species.

Here, we developed the Grass Expression Atlas (GExA), an interactive and user-friendly web-based gene expression database for comparative exploration of public RNA-seq data in grasses. GExA integrates publicly available RNA-seq data for four millet species—pearl millet, foxtail millet, proso millet (*Panicum miliaceum*), and finger millet (*Eleusine coracana*)—together with barley (*Hordeum vulgare*) and sorghum (*Sorghum bicolor*) as reference species. Importantly, the code underlying GExA was designed with generality and automated updating in mind, with the aim of overcoming both the lack of a practical pearl millet RNA-seq database and the limited usability and comparability of existing millet resources. By integrating public RNA-seq datasets through a unified workflow and implementing a user-friendly web interface, GExA aims to improve the reusability of public transcriptome data in millet research and to provide a foundation that accelerates candidate gene discovery and hypothesis generation.

## Results

### RNA-seq datasets

GExA hosts RNA-seq datasets for six plant species: pearl millet, foxtail millet, proso millet, finger millet, barley, and sorghum. The database comprises 987 pearl millet samples from 43 BioProjects, 2,216 foxtail millet samples from 211 BioProjects, 117 finger millet samples from 20 BioProjects, 737 proso millet samples from 27 BioProjects, 345 barley samples from 27 BioProjects, and 271 sorghum samples from 114 BioProjects (Table 1). For pearl millet, we used genome assemblies from three cultivars (Tift23D_2_B_1_-P1-P5 (Tift), ICMB843, and ICMR06777) as reference genomes (Ramu et al. 2023), enabling cross-cultivar comparisons of gene expression. All datasets were retrieved and reanalyzed using the standardized processing framework summarized in Fig. 1. The workflow has been fully scripted, and the source code and usage instructions are publicly available via a GitHub repository (https://github.com/kotakambara/GExA-pipeline).

**Table 1.** Summary of RNA-seq datasets and reference genomes included in GExA.

|  | Number of<br>samples | Number of<br>projects | Reference | Cultivar |
| --- | --- | --- | --- | --- |
| Pearl millet | 987 | 43 | Ramu et al. 2023 | Tift, ICMB843, and<br>ICMR06777 |
| Foxtail millet | 2216 | 211 | Bennetzen et al. 2012 | Yugu1 |
| Finger millet | 117 | 20 | Devos et al. 2023 | KNE 796-S |
| Proso millet | 737 | 27 | Wang et al. 2024 | AJ8 |
| Barley | 345 | 27 | Navrátilová et al. 2022 | Morex |
| Sorghum | 271 | 114 | Paterson et al. 2009 | BTx623 |

**Fig. 1.**
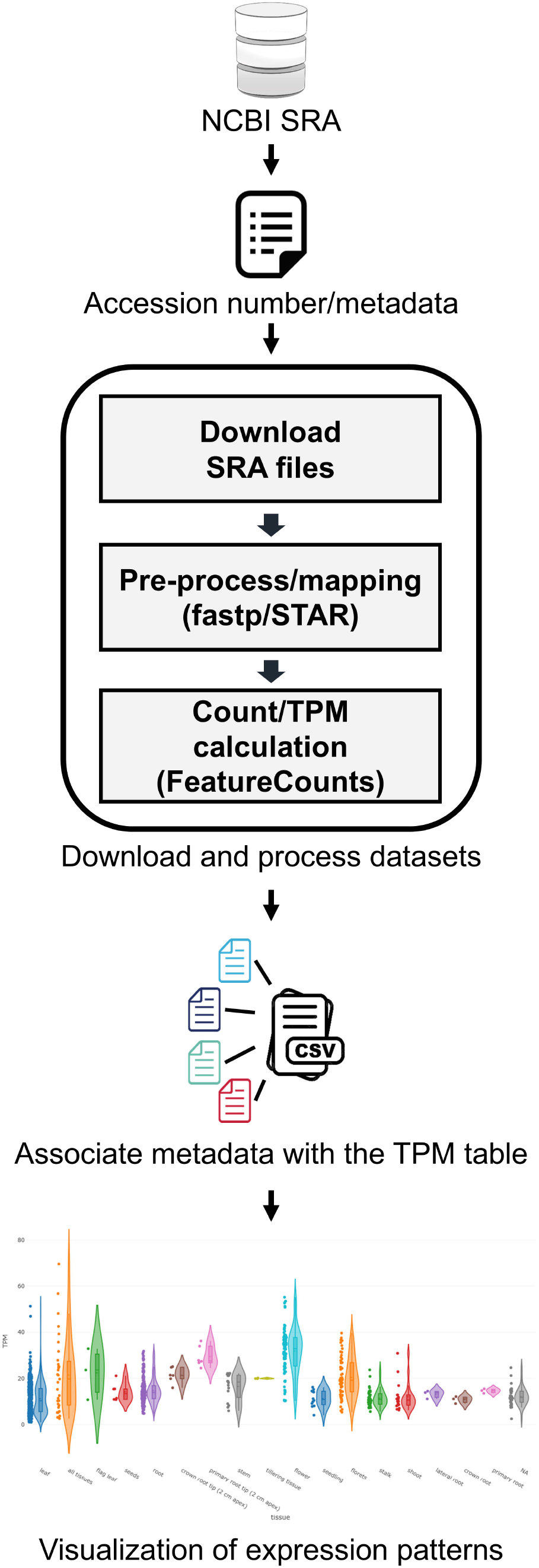
Workflow for data collection and processing to construct GExA. Public RNA-seq raw reads and associated sample metadata were retrieved from the National Center for Biotechnology Information (NCBI) Sequence Read Archive (SRA). Datasets were processed using a pipeline consisting of read preprocessing and quality filtering (fastp), genome alignment (STAR or HISAT2, depending on species), and gene-level quantification (featureCounts), followed by conversion to transcripts per million (TPM). The resulting TPM table was integrated with curated sample metadata and deployed for interactive visualization of expression patterns in the GExA web interface. Alt text: Vertical flowchart summarizing construction of the GExA expression matrix. Public RNA-seq data are sourced from the NCBI Sequence Read Archive (SRA) and linked to accession numbers and sample metadata. A boxed processing pipeline shows sequential steps—downloading SRA files, read preprocessing/mapping, and gene counting with TPM calculation—followed by integration of curated metadata with the resulting TPM table and downstream visualization of expression patterns.

Principal component analysis (PCA) was performed for each species, and the distributions of samples on PC1 and PC2 are shown in Fig. 2 and Supplementary Fig. S1–S3. In these plots, samples were separated according to tissue or organ type, suggesting that the processed expression profiles reflected major biological differences among samples.

**Fig. 2.**
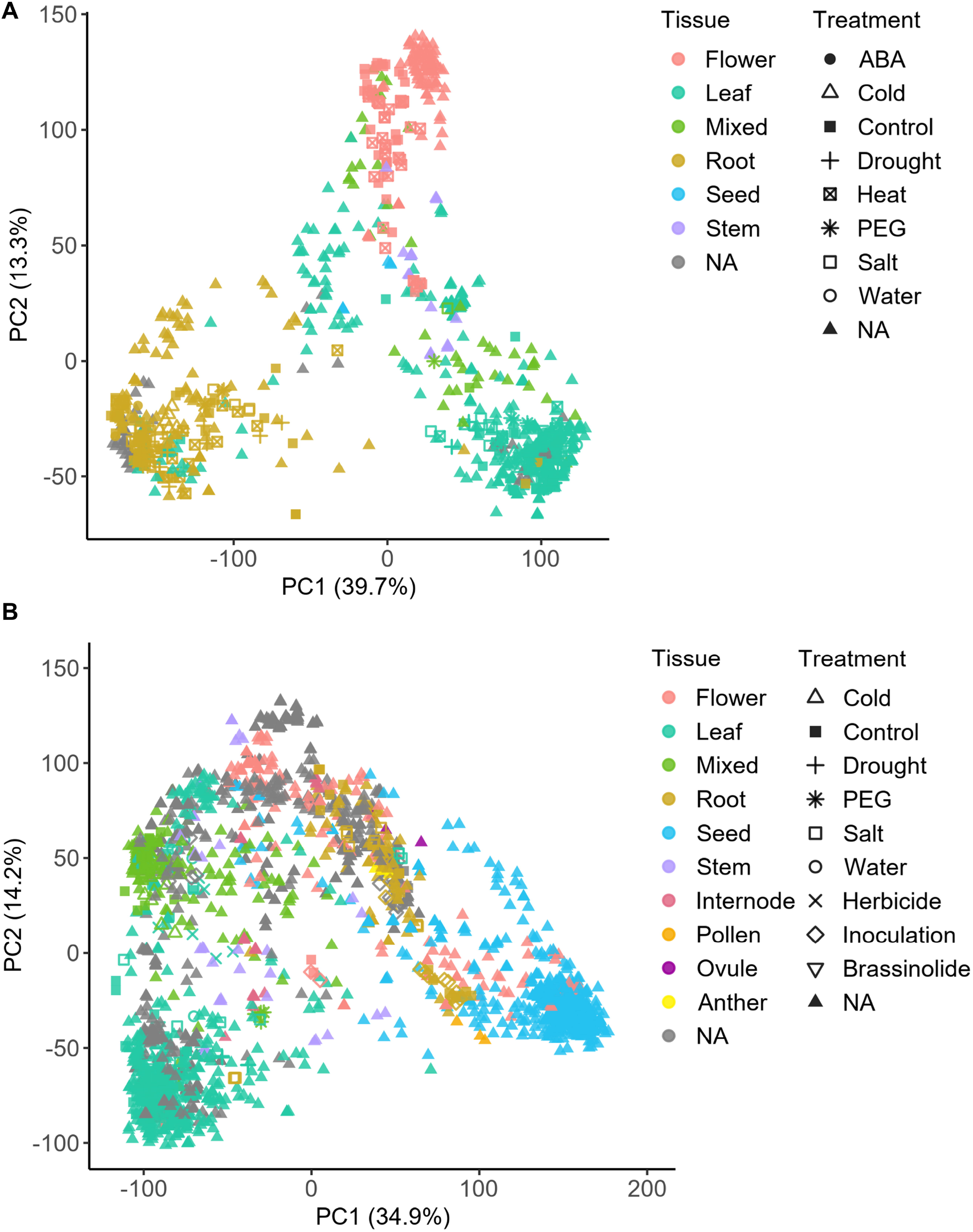
Principal component analysis of RNA-seq samples in pearl millet and foxtail millet. PCA was performed using TPM values from 987 pearl millet samples mapped to the reference genome of cultivar Tift (A) and 2,216 foxtail millet samples (B), based on the 5,000 most variable genes after filtering lowly expressed genes (TPM < 1). Each point represents one RNA sequencing (RNA-seq) sample. Points are colored by the curated “tissue” category and shaped by a manually curated treatment category with modification. The percentages of variance explained by PC1 and PC2 are shown on the respective axes. Alt text: Two principal component analysis (PCA) scatterplots of RNA-seq samples based on TPM values from the most variable genes: (A) pearl millet and (B) foxtail millet. Each point represents one RNA-seq sample positioned by PC1 and PC2 (with percent variance explained shown on the axes). Points are color-coded by curated tissue categories and use different marker shapes to indicate curated treatment categories; legends for tissue and treatment are shown to the right of each panel.

### Web interface

GExA is provided as an interactive web application for querying and visualizing gene expression patterns. Users first select a species from the main menu, which opens the corresponding search page (Fig. 3A).

**Fig. 3.**
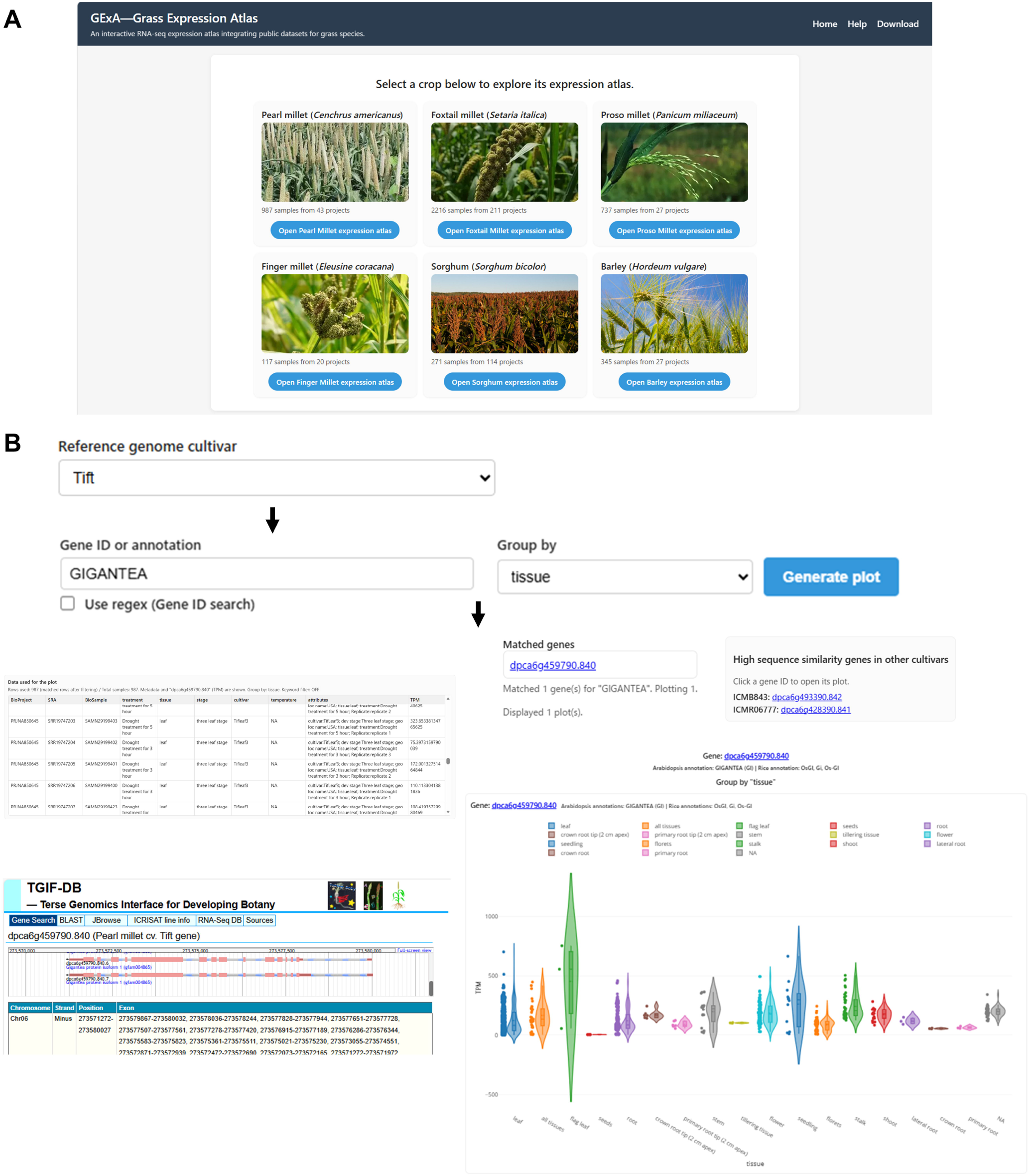
GExA web interface and representative query functions. (A) The GExA main page, showing species selection for six crops (pearl millet, foxtail millet, proso millet, finger millet, sorghum, and barley). (B) Example workflow for keyword-based search in pearl millet. Users select a cultivar (e.g., Tift), enter a gene ID or an annotation keyword (e.g., “GIGANTEA”), choose a metadata category for grouping (e.g., tissue), and generate an expression distribution plot. Matched genes are listed, and highly similar genes in the other reference genomes are provided with direct links for cross-cultivar comparison. The “Data used for the plot” table reports per-sample TPM values together with associated metadata. Gene IDs displayed above the plot link to the corresponding TGIF-DB entries (Tsugama and Takano, 2021). Alt text: Screenshots illustrating the GExA web interface and a typical query workflow. (A) The main page presents six crop tiles (pearl millet, foxtail millet, proso millet, finger millet, sorghum, and barley), each with an image and a button to open the corresponding expression atlas. (B) The query page for pearl millet shows a cultivar selector (set to “Tift”), a text field for gene ID or annotation keyword (example: “GIGANTEA”), a “group by” selector (example: tissue), and a “Generate plot” button, with arrows indicating the stepwise workflow. Output sections include a list of matched genes, links to highly similar genes in other cultivars for cross-reference, a per-sample data table (TPM values with associated metadata), and an expression distribution plot; an example external gene-entry page (TGIF-DB) is also shown.

One or more gene IDs (optionally specified using regular expressions) can be entered to visualize transcripts per million (TPM) distributions using jittered dot plots or violin plots. Alternatively, users can search by keywords (e.g. a protein name such as “GIGANTEA”); keyword searches return genes whose homology-based annotations, derived from BLASTP searches against Arabidopsis or rice proteins, contain the queried term. (Fig. 3B). Samples can be grouped by BioProject, tissue, treatment, developmental stage, temperature, or cultivar, and can be filtered using free-text queries over sample metadata. For pearl millet datasets represented by reference genomes from three cultivars, the interface provides a reference-genome selector that directs users to genome-specific pages. When the cultivar “Tift” is selected, a similarity panel lists gene IDs of high sequence-similarity genes in the other cultivars (ICMB843 and ICMR06777) and provides direct links to the corresponding expression pages, enabling rapid cross-cultivar comparisons. Clicking a gene ID displayed above the plot redirects the user to the corresponding TGIF-DB entry (Tsugama and Takano 2021) for detailed gene information (Fig. 3B). Each plot is accompanied by a “Data used for the plot” table listing per-sample TPM values together with the associated metadata. This enables users to identify the contributing BioProjects and sample attributes (e.g. tissue, developmental stage, and cultivar) in each group and to interpret expression patterns across diverse experimental designs (Fig. 3B).

### Example application

To illustrate the utility of GExA, we examined one pearl millet gene dpca1g022930.840 (corresponding to Pgl_GLEAN_10033150 in a genome assembly in Varshney et al. 2017). This gene is the closest homolog of rice *HVA22* (LOC_Os11g30500) and was recently reported as a dehydration- and abscisic acid (ABA)-responsive gene in pearl millet (Qazi et al. 2025).

Using the pearl millet page, users select the Tift reference genome and enter dpca1g022930.840 in the search box. By selecting “treatment” in the “*Group by*” menu and generating the plot, the expression distribution is visualized across treatment groups. In this query, dpca1g022930.840 shows higher TPM values in multiple drought- and ABA-related treatment groups (Fig. 4), consistent with previous findings on drought/ABA inducibility (Qazi et al. 2025).

**Fig. 4.**
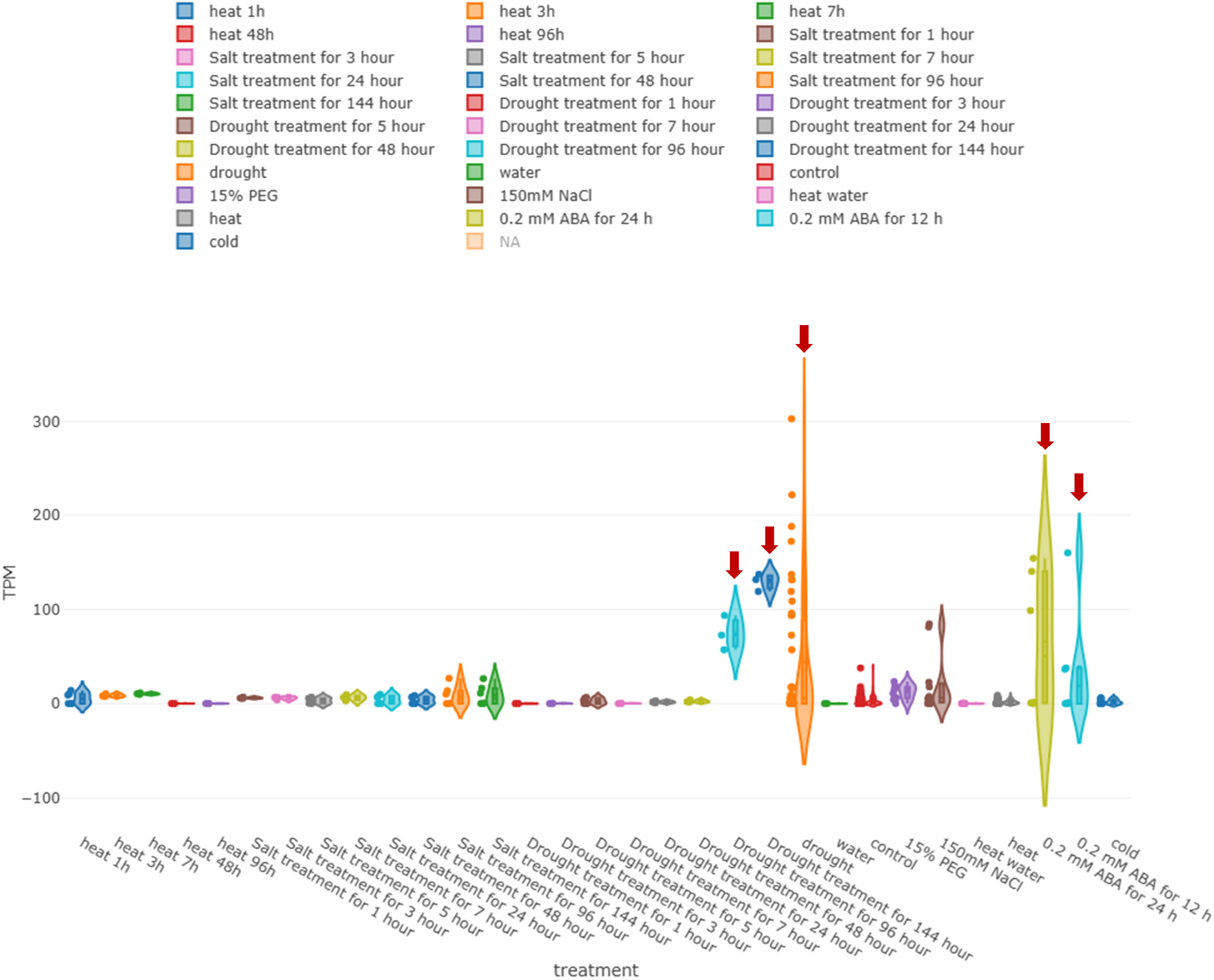
Example application of GExA: expression of a pearl millet *HVA22*-like gene across treatments. An expression distribution plot was generated for the pearl millet gene dpca1g022930.840, grouped by treatment. The x-axis indicates treatment categories and the y-axis indicates TPM values. Each dot represents one RNA sequencing (RNA-seq) sample. Red arrows highlight drought- and abscisic acid (ABA)-related treatment groups exhibiting higher TPM values. Alt text: Expression distribution plot for the pearl millet *HVA22*-like gene dpca1g022930.840 grouped by treatment. The x-axis lists multiple treatment categories, and the y-axis shows expression in TPM. For each treatment, a distribution shape is overlaid with individual sample points to show both spread and sample-level values. Red arrows mark selected drought-related and abscisic acid (ABA)-related treatment groups that display higher expression compared with most other treatments.

## Discussion

GExA provides an integrated and user-friendly transcriptome resource for millet species and related grasses, facilitating comparative exploration of gene expression across tissues, developmental stages, and experimental conditions. GExA addresses several limitations of previously available millet transcriptome resources. In our previous study, public pearl millet RNA-seq datasets were collected, reanalyzed, and released as expression matrices (Kambara et al. 2024), but that resource relied on an earlier reference genome assembly (Varshney et al. 2017) and did not provide a web interface for interactive exploration. In the present study, we updated the analysis using higher-quality pearl millet reference genome assemblies published in 2023 (Ramu et al. 2023), expanded the dataset to 987 samples, and implemented an interactive web interface that enables users to group and compare expression profiles according to tissue, developmental stage, treatment, cultivar, and other metadata categories. These improvements substantially enhance the accessibility and reusability of public transcriptome data for pearl millet research.

A limitation of transcriptome analyses that rely on a single reference genome is reference bias, where sequence divergence between a sample and the reference can affect read mapping and downstream quantification (Chen et al. 2021). To mitigate this issue, GExA adopts a multi-reference design for pearl millet by providing expression pages based on three reference genomes (Tift, ICMB843, and ICMR06777). This design allows users to examine whether observed expression patterns are robust to the choice of reference genome and thus helps address potential quantification biases associated with reference selection (Chen et al. 2021).

Another key feature of GExA is its scalability. The pipeline developed in this study automates data retrieval, read mapping, and downstream processing (Fig. 1), and the source code is publicly available (https://github.com/kotakambara/GExA-pipeline). Consequently, the workflow can be applied to additional species for which reference genome sequences are available. In future work, we plan to extend the framework beyond the four millet species currently included and to expand GExA to additional plant species.

Despite these advantages, GExA is based exclusively on publicly available data and therefore has limitations. Integrating datasets across independent projects inevitably introduces heterogeneity arising from differences in library preparation, sequencing platforms and parameters, growth conditions, and experimental designs, which can lead to batch effects and confounding (Leek et al., 2010). In GExA, samples were processed using a standardized processing framework and transcript abundance was quantified using TPM to reduce variability attributable to analytical procedures. Nevertheless, such processing cannot fully correct for biological and technical differences across studies, and observed expression differences may not necessarily reflect true biological effects. Therefore, GExA is primarily intended as a foundation for exploratory analyses and hypothesis generation. For rigorous differential expression testing or causal interpretation, findings should be validated using independent datasets generated under appropriately controlled conditions.

In summary, we developed the Grass Expression Atlas (GExA), a user-friendly, web-based, interactive gene expression database. GExA systematically collects and uniformly processes public RNA-seq data for four millets (pearl millet, foxtail millet, proso millet, and finger millet), with barley and sorghum included as reference species. GExA enables rapid visualization of transcript abundance distributions across tissues, developmental stages, and experimental conditions, and provides access to per-sample TPM values together with curated metadata. By improving the accessibility and reusability of public RNA-seq data, GExA provides a foundation for exploratory analyses and hypothesis generation in these crops.

## Materials and Methods

### Data collection

For pearl millet, foxtail millet, proso millet, and finger millet, publicly available RNA-seq datasets were identified in the National Center for Biotechnology Information (NCBI) Sequence Read Archive (SRA) as of December 2025. Candidate records were retrieved using the NCBI Entrez Programming Utilities (E-utilities) with a query combining organism terms and transcriptome-related keywords (e.g., “RNA-seq” OR “transcriptome”). Queries and record retrieval were automated using a custom Python script (API_RNA-seq.py). From the returned SRA summaries, we generated a sample list containing the run accession, BioSample accession, BioProject accession, sample name, and library layout (paired-end or single-end). This list served as the primary input for the mapping and quantification pipeline (Mapping_script.sh). Sample metadata were extracted from BioSample records, including cultivar, tissue, treatment, developmental stage and temperature, where available. For barley (*Hordeum vulgare*) and sorghum (*Sorghum bicolor*), we used a curated list of datasets summarized in Supplementary Table S1.

### Processing of RNA-seq data

Raw sequencing files were downloaded using aria2c from the NCBI SRA public cloud repository. For runs archived in the DNA Data Bank of Japan (DDBJ) Sequence Read Archive (DRA), data were retrieved from the DDBJ public repository. Downloaded SRA files were converted to FASTQ format using the fasterq-dump command in SRA Toolkit v3.3.0 (https://www.ncbi.nlm.nih.gov/sra/docs/toolkitsoft/). To avoid inclusion of extremely small datasets, samples with a total raw FASTQ size of <100 MB were excluded. Adapter trimming and quality filtering were performed using fastp v0.23.4 (Chen 2023) with default parameters. For pearl millet, proso millet, finger millet and foxtail millet, reads were mapped to the corresponding reference genome assemblies (Table 1) using STAR v2.7.11 (Dobin et al. 2013) with default parameters. For barley and sorghum, reads were mapped to the corresponding reference genome assemblies (Table 1) using HISAT2 v2.2.1 (Kim et al. 2019) with default parameters. Gene-level read counts were obtained using featureCounts v2.0.6 (Liao et al. 2014) with the corresponding GFF3 annotation files. Obtained read counts were converted to transcripts per million (TPM) using a custom Python script (Count_to_TPM.py). The resulting TPM tables were merged with sample metadata, including BioProject ID, Run accession numbers, BioSample ID, treatment, tissue, stage, cultivar, sample name, temperature and additional metadata fields. To improve the consistency of metadata in grouping and filtering within GExA, key metadata fields (notably tissue, treatment, developmental stage, and cultivar) were harmonized to reduce orthographic variation using a custom Python script (standardize_metadata_XX.py). To remove low-quality samples, we excluded libraries in which >90% of annotated genes had TPM <0.01, using a custom Python script (check_sample.py). The final tables containing sample attributes, read counts, and TPM values were deposited in the figshare repository at https://doi.org/10.6084/m9.figshare.31669273.

### Gene annotation based on Arabidopsis and rice

Protein-coding genes of each target crop species were functionally annotated by homology-based association with Arabidopsis and rice proteins, which represent well-annotated dicot and monocot model species, respectively, following the approach described in Tsugama and Takano (2021). Protein sequences and functional annotations for Arabidopsis (TAIR10) were obtained from The Arabidopsis Information Resource (TAIR) (Berardini et al. 2015). Protein sequences and annotations for rice were obtained from the Rice Genome Annotation Project (Kawahara et al. 2013) and Oryzabase (Kurata and Yamazaki 2006). BLASTP searches were performed using BLAST+ (Camacho et al. 2009). Protein sequences from each crop species were used as queries against either the Arabidopsis or rice protein datasets as the search database. Hits with an E-value <1e-20 were retained, and each crop protein was assigned to the functional annotations of the corresponding Arabidopsis and/or rice proteins.

### Web service implementation

The web service was implemented using HTML, CSS, and JavaScript. Interactive visualizations are rendered in the user’s web browser using Plotly.js. To maintain responsiveness when handling large-scale expression matrices, the gene expression matrix was converted into web-optimized static resources with a custom Python script (preprocess_shard.py). Sample metadata are provided as a tab-delimited file (meta.tsv), while a gene index (gene_index.tsv) and a manifest (manifest.json) map each gene to its corresponding binary shard and column position. The expression values are stored in multiple binary shard files, enabling the client to fetch only the minimal subset of data required for a given query. The website is hosted on a rental server provided by SAKURA Internet Inc. (Osaka, Japan).

## Data and Code Availability

All the data and codes are available in the figshare repository at https://doi.org/10.6084/m9.figshare.31669273.

## Acknowledgments

The authors thank our colleagues for their assistance in testing earlier versions of GExA. The authors used ChatGPT (OpenAI) for code generation and debugging during the development of the expression atlas. The authors have verified the accuracy and functionality of all AI-generated code and take full responsibility for the results.

## Author Contributions

All authors contributed to designing the study. KK and DT performed data gathering and processing. KK and QC constructed the database and developed the website. DT supervised the study. KK and DT wrote the manuscript, and all authors critically revised the manuscript and gave final approval.

