## Supplementary Figures S1-S3 for "Grass Expression Atlas: an RNA-seq-based expression resource for grass species"

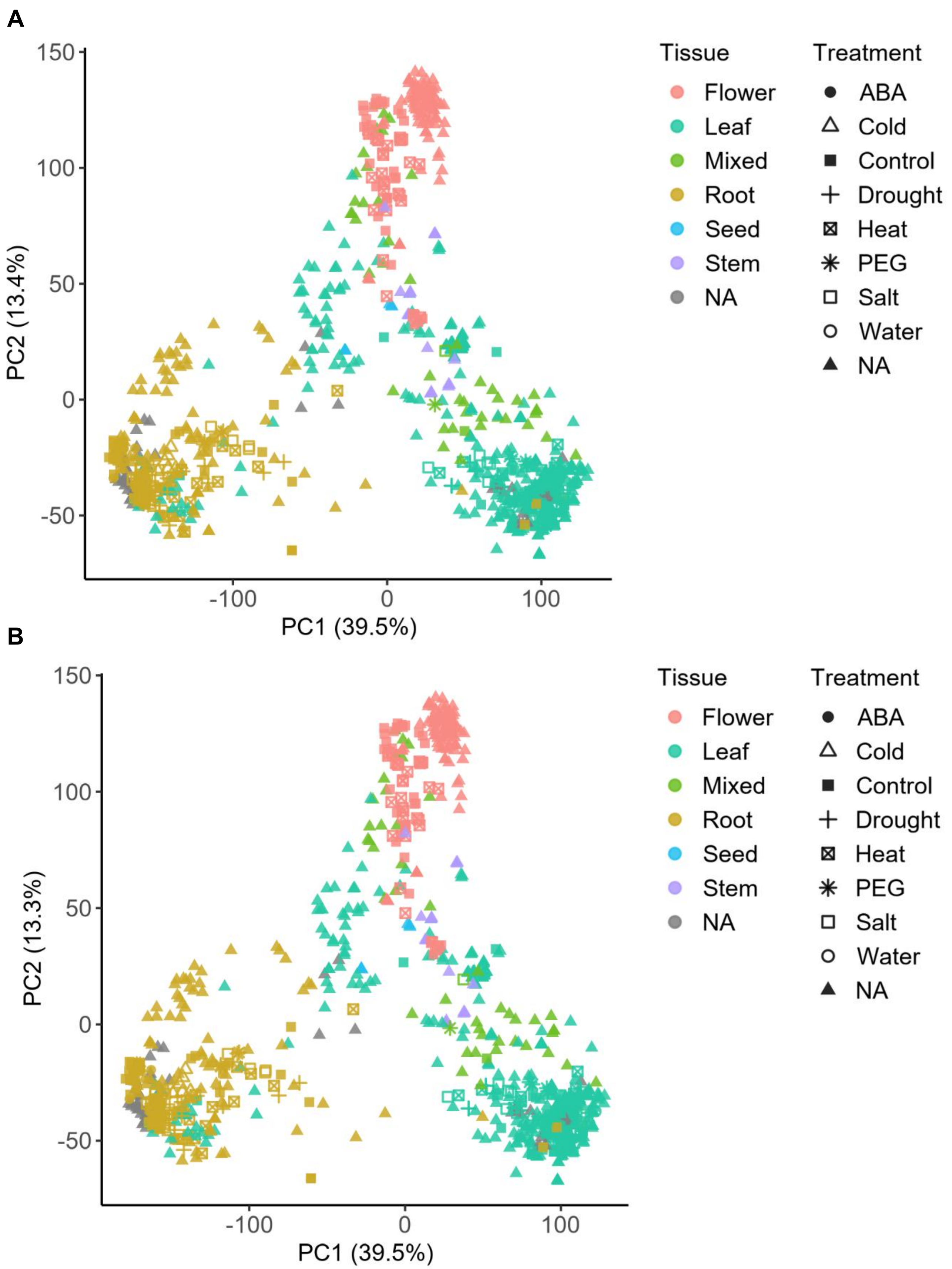


Supplementary Fig. S1. Principal component analysis of RNA-seq samples in pearl millet. PCA was performed using TPM values from 987 pearl millet samples mapped to reference genome cultivar "ICMB843" (A) and "ICMR06777" (B), based on the 5,000 most variable genes after filtering lowly expressed genes (TPM < 1). Each point represents one RNA sequencing (RNA-seq) sample. Points are colored by the curated "tissue" category and shaped by a manually curated treatment category with modification.The percentages of variance explained by PC1 and PC2 are shown on the respective axes.

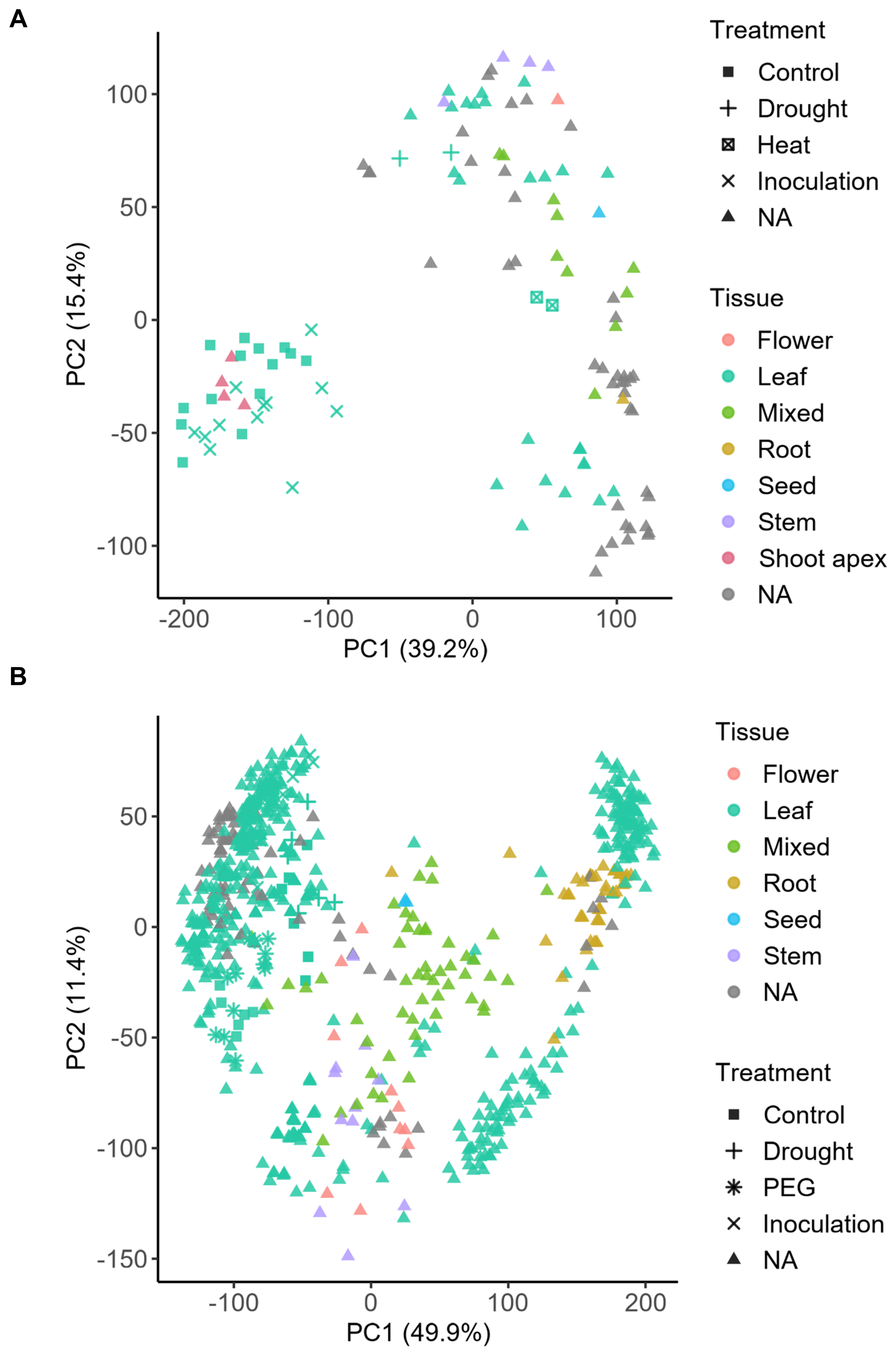


Supplementary Fig. S2. Principal component analysis of RNA-seq samples in finger millet and proso millet. PCA was performed using the same procedure as in Supplementary Fig. S1, using TPM values from 117 finger millet samples (A) and 737 proso millet samples (B), based on the 5,000 most variable genes after filtering lowly expressed genes (TPM < 1).

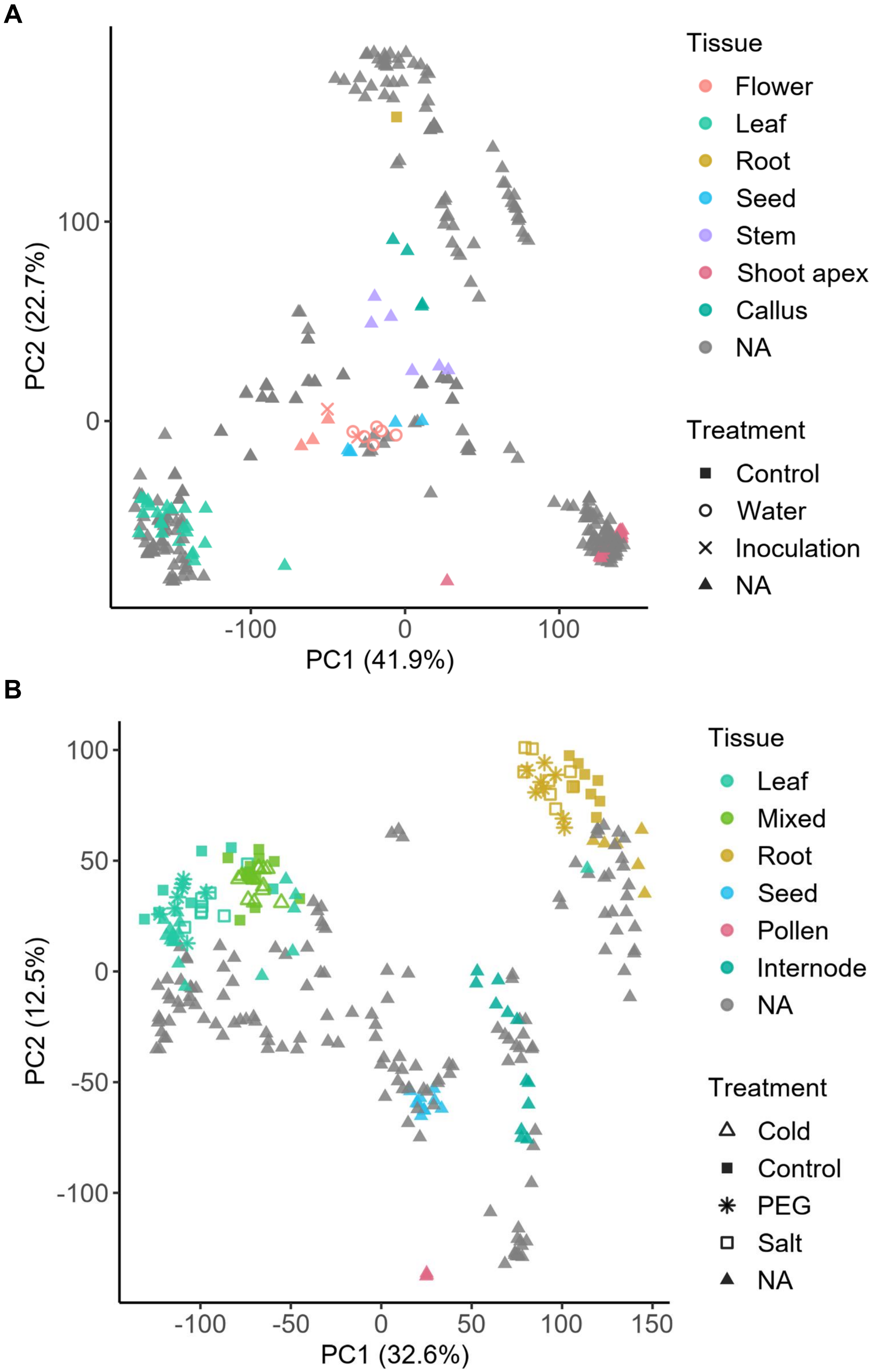


Supplementary Fig. S3. Principal component analysis of RNA-seq samples in barley and sorghum. PCA was performed using the same procedure as in Supplementary Fig. S1, using TPM values from 345 barley samples (A) and 271 sorghum samples (B), based on the 5,000 most variable genes after filtering lowly expressed genes (TPM < 1).
